# A Double-Blind Replication Attempt of Offline 5Hz-rTUS-Induced Corticospinal Excitability

**DOI:** 10.1101/2024.11.25.625187

**Authors:** Po-Yu Fong, Benjamin R. Kop, Carys Evans, Vidya Gani Wijaya, Yongling Lin, Drew Cappotto, Jenny S. A. Lee, Anna Latorre, Joy Song, Bradley Treeby, Eleanor Martin, John Rothwell, Lennart Verhagen, Sven Bestmann

## Abstract

**Introduction:** Transcranial ultrasound stimulation (TUS) is a promising new form of non-invasive neuromodulation. As a nascent technique, replication of its effects on brain function is important. Of particular interest is offline 5Hz repetitive TUS (5Hz-rTUS), originally reported by Zeng and colleagues [1] to elicit lasting corticospinal excitability increases, with large effect sizes.

**Material and method:** Here, we conducted a pre-registered (https://osf.io/p5n4q) replication of this protocol that benefitted from three additional features: double-blind application of TUS, neuronavigation for consistent TMS positioning, and acoustic simulations to assess M1 target exposure to TUS. Changes in resting motor thresholds (rMT), motor-evoked potential (MEP) amplitude, short-interval intracortical inhibition (SICI) and intracortical facilitation (ICF) in response to TUS (5Hz-rTUS vs. sham) were measured in the right first dorsal interosseous (FDI), abductor digiti minimi (ADM) and abductor pollicis brevis (APB) muscles, with unbiased selection of participants. Transducer location was determined by the TMS-hotspot for motor representations of the right FDI, as in the original work.

**Results:** No significant effects of 5Hz-TUS (vs sham) were observed. Post-hoc simulations showed considerable variability of the acoustic focus, which was outside the anatomical M1-hand area in 67% of participants – in line with the known poor correspondence of TMS-hotspot location and M1-hand area.

**Conclusion:** Our results indicate that the effect sizes of the neuromodulatory effects of 5Hz-rTUS on M1 may be more variable than previously appreciated. We suggest that double-blinding, neuronavigated TMS, individualised acoustic simulations for TUS targeting and pre-registration will aid reproducibility across studies.

## Introduction

Transcranial ultrasonic stimulation (TUS) is a relatively novel technique for non-invasive neuromodulation in humans [2]. As a nascent approach for neuromodulation, replication of the effects of TUS is a critical ingredient in progressing toward a mature technology with clinical utility. One example illustrating this is the recent discovery that the inhibitory effects of TUS on corticospinal excitability can be confounded by auditory inputs [3]. In response, auditory masking and ramped pulsing have rapidly become standard in the field [4–7]

Here we focus on 5Hz-rTUS (also referred to as theta-burst TUS; tbTUS), an offline TUS protocol that has been reported to elicit strong facilitation of corticospinal excitability (CSE), with large effects sizes, that outlasted sonication by up to 30 minutes [1]. The same research group has subsequently replicated these excitatory effects in both healthy and clinical populations [8–14]

However, the opposite neuromodulatory effects of 5Hz-rTUS were recently reported by an independent research group, instead showing inhibition (not excitation) of CSE lasting up to 30 minutes following sonication [15]. In contrast with targeting M1 based on the TMS-hotspot location and only at a fixed depth of ∼30mm [1], Bao and colleagues personalised TUS application with acoustic simulations to ensure precise targeting within M1 in each participant. The inhibitory effects were observed when sonicating either the lip of precentral gyrus or deeper sections of M1. Simulations also provided individual estimates of the acoustic intensity in the brain, as opposed to using measurements in free-water which cannot account for individual variance in absorption and refraction of ultrasound through the skull [16, 17]. Given the contrasting outcomes of 5Hz-rTUS reported by separate research groups, and a dearth of independent replication, at this early stage of offline human TUS application it is essential to obtain better estimates of the likely effect sizes of 5Hz-rTUS and factors that may influence these. This necessity is further underscored by the intended application of these protocols in clinical populations [12].

In the present study, we therefore conducted an independent replication of the methodology initially reported by Zeng and colleagues [1]. We specifically focussed on replicating the effect of 5Hz-rTUS on CSE, short-interval intracortical inhibition (SICI), and intracortical facilitation (ICF). We employed the same targeting procedures, stimulation protocol, and outcome measures as [1]. However, to guard against biases and additional sources of variance, we incorporated double-blinding, neuronavigation of TMS, more repetitions per condition, quantification of MEPs from adjacent hand muscles, post-hoc acoustic simulations of TUS, and pre-registered the study.

We did not observe a significant effect of 5Hz-rTUS on CSE or intracortical excitability. This result reappraises the likely effect sizes of 5Hz-rTUS, and we highlight potential factors influencing these discrepancies.

## Materials and methods

Unless stated otherwise, all elements of Experiment 2 by Zeng and colleagues [1] were adhered to.

### Design

Participants attended two sessions (one week apart) with either 5Hz-rTUS or sham-TUS, in counterbalanced order across participants (Fig. 1A-C). In addition to Zeng and co-workers [1], we used a double-blind procedure for TUS application to reduce potential experimenter bias. MEP measures (MEP amplitude; SICI; ICF) were recorded at baseline, and 5, 30, and 60 minutes after 5Hz-rTUS or sham (T5, T30, T60; Fig 1D). Each session was conducted by two experimenters – one operating the TMS and TUS, and the other assisting with neuronavigation and stimulation equipment control.

**Fig. 1.**
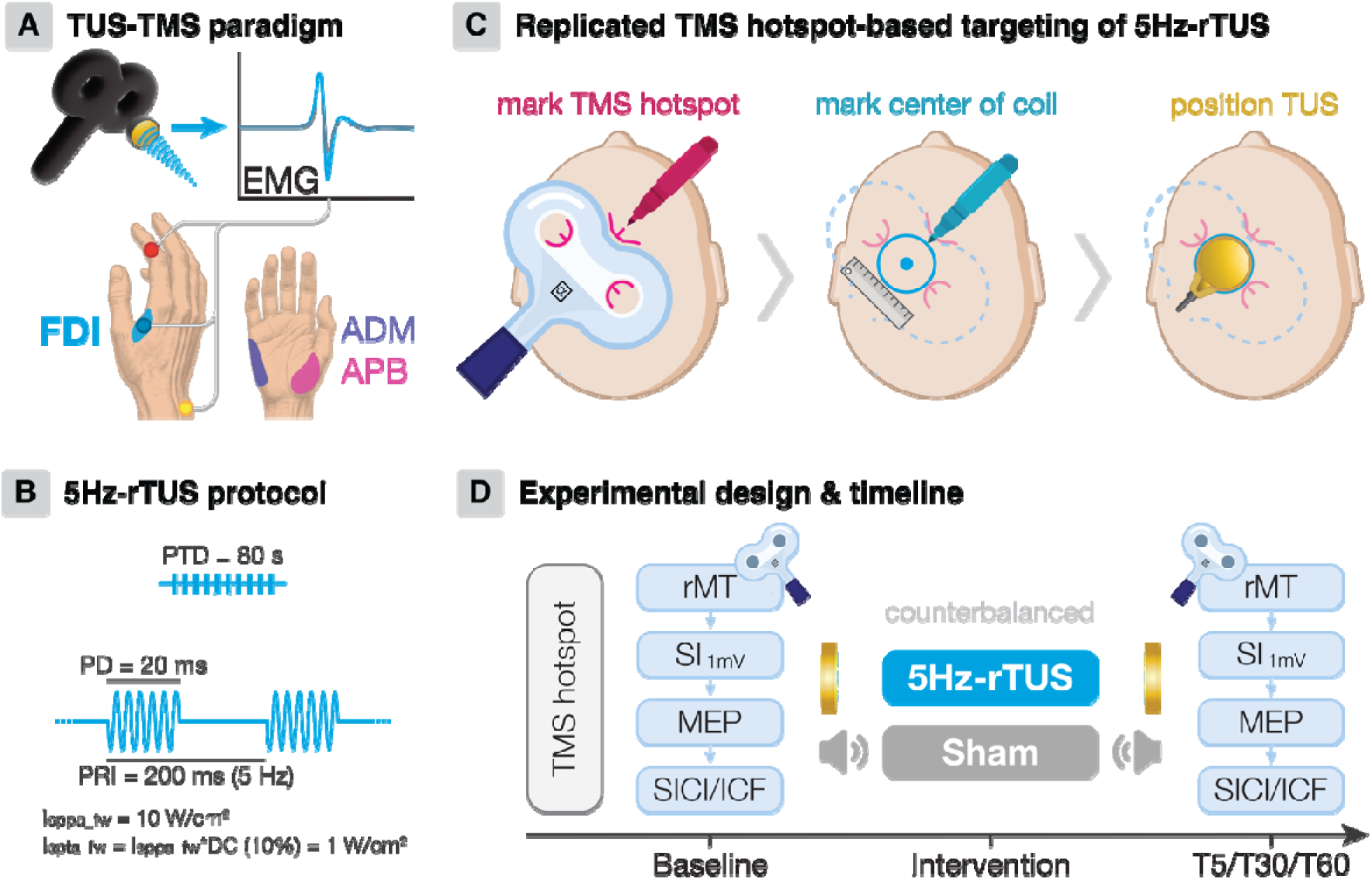
Experimental design and methodology. **A.** TMS-elicited motor-evoked potentials (MEPs) were measured from the primary muscle of interest (FDI) and two adjacent hand muscles. **B.** The 5Hz-rTUS protocol. PTD: pulse train duration; PD: pulse duration; PRI: pulse repetition interval; DC: duty cycle; f I_sppa_fw_: free-water spatial-peak pulse-average intensity; I_spta_fw_: free-water temporal-average spatial peak. **C.** TUS positioning [1]: TMS coil position over the motor hotspot was marked on the scalp (left), with the centre of the coil (middle) used to localise the TUS transducer on the scalp (right). **D.** Experimental design and procedure.

### Participants

We studied 15 healthy right-handed individuals (age: 31.3±12; five males; eight Asian, six Caucasian, one African) after written informed consent, all without neurological or psychiatric diseases, no contraindications to brain stimulation, and no medications known to affect brain excitability. The study was approved by the UCL Research Ethics Committee (14233/003) and conducted in accordance with the Declaration of Helsinki.

### MRI

All participants had existing T1-weighted MRI scans obtained in one of UCL’s neuroimaging facilities, and all consented to re-use of these images for this study. While imaging sequence details varied for this reason, all images were acquired with 1mm isotropic resolution, covering the whole head. Note that for the simulation conducted in this study, we focus on the location of the acoustic focus, not the actual intensity values, which will not have been systematically influenced by small idiosyncrasies in the MRI sequences across individuals.

### Transcranial magnetic stimulation

TMS was delivered with a Magstim 200^2^ Monophasic stimulator (single-pulse) and a Magstim Bistim^2^ system consisting of two 200^2^ monophasic stimulators (paired-pulse) connected to a D70 alpha F8 coil with an internal diameter of 70mm (Magstim Co. Ltd, Whitland, Wales). Participants were seated in a comfortable chair with their hands resting on a pillow on their lap. The TMS coil was placed over the left primary motor cortex (M1), tangentially to the scalp, with the handle of the coil pointing backwards at a 45-degree to the midline to induce an approximate posterior-anterior current. The TMS-hotspot was defined as the area of the scalp where the largest and most stable MEPs were observed in the right first dorsal interosseus (FDI) muscle [18] (Fig. 1A). Apart from the setup in the original work [1], neuronavigation was used to ensure precise and consistent placement of TMS both before and after 5Hz-rTUS/sham (Brainsight, Rogue Research, USA). We additionally recorded MEPs from adjacent abductor pollicis brevis (APB) and abductor digiti minimi (ADM) muscles, to capture any potential effects of the ultrasound beam on other intrinsic hand muscles (Fig. 1A).

Our TMS procedures followed the original study [1]. Briefly, after identifying the TMS hand motor hotspot, we determined the baseline rMT and the stimulator intensity (SI) required to elicit a ∼1mV MEP (SI_1mV_) [18, 19]. This baseline SI_1mV_ was the intensity for all subsequent single-pulse MEP blocks to assess changes in CSE. For the paired-pulse TMS block capturing intracortical excitability, the intensity of the conditioning stimulus was set to 80% rMT for SICI (2 ms ISI) and ICF (10 ms ISI), and the test stimulus was set to the SI_1mV_ measured in the beginning of the same block. Paired-pulse trials were administered in pseudorandomized order. In contrast to Zeng and co-workers’ work [1], we acquired 25 trials for each measurement – originally 20 and 10 trials for each single- and paired-pulse metric, respectively – using an ISI of 5 seconds and a 10% jitter. All TMS measures were acquired at baseline, T5, T30 and T60 (Fig. 1D). Each TMS block lasted approximately nine minutes.

Surface electromyography (EMG) signals were simultaneously recorded for target FDI, APB and ADM with pairs of surface electrodes in a belly-tendon montage (WhiteSensor 40713, AmbuR, Denmark; Fig. 1A). Signals were amplified with a gain of 1000, bandpass filtered (5 Hz - 3000 Hz) by a Digitimer D360 amplifier (Digitimer Ltd, Welwyn Garden City, Hers, UK), and digitised at 5000 Hz by a Power 1401 data acquisition interface and Signal software version 7.01 (Cambridge Electronic Design Ltd., Cambridge, UK).

### Transcranial ultrasound stimulation

Transcranial ultrasound stimulation (TUS) was delivered using the NeuroFUS system (manufacturer: Sonic Concepts Inc., Bothell, WA, USA; supplier/support: Brainbox Ltd., Cardiff, UK) via a four-element 500 kHz annular array piezoelectric transducer with a 64 mm radius of curvature and aperture diameter (CTX-500-025, Sonic Concepts Inc., Bothell, WA, USA). A four-channel radiofrequency amplifier (Transducer Power Output system; TPO) powered the transducer via a four-channel electrical impedance matching network utilising a rectangular pulse shape. The transducer was coupled with a 10 mm gel pad (Aquaflex, Parker, Laboratories, NJ, USA). Ultrasound gel (Aquasonic 100, Parker Laboratories, NJ, USA) was centrifuged to remove visible bubbles and applied between the gel pad and the transducer. We defined our sonication depth as 33 mm in accordance with the acoustic field peak reported previously [1, 12]. To reach 33 mm, the sonication depth on the TPO was set at 43.5 mm to account for the additional distance added by ultrasound gel (0.5mm) and gel pad (10mm). Table S1 and Figure S1 show the comparison of the acoustic intensities and profiles between Zeng et al. (2022) and the current study. The applied focal depth in the present study and the original study [1] matches the scalp-to-cortex distance of ∼30mm for the M1 omega formation.

The 5Hz-rTUS protocol was an 80-second train of 20-millisecond ultrasound pulses repeated every 200 milliseconds (PRF 5Hz; 10% duty cycle), for a total of 400 pulses (Fig. 1B). The spatial-peak-pulse average intensity in free water (I_SPPA_) was set to 10 W/cm^2^ (Table S1), similar to the original work. Applying an estimated 75% skull attenuation relative to free-water measurements, our estimated transcranial I_SPPA_ (2.5W/cm^2^) was consistent with the original study (2.26W/cm^2^) [1]. Our actual estimated transcranial I_SPPA_ based on acoustic simulations was 1.20W/cm^2^ ± 0.43.

The TUS transducer location was determined by the TMS-hotspot, as previously [1, 9–11, 13, 14]. The contours of the TMS coil were marked on the scalp using a chinagraph pencil and the centre of the TMS coil was measured (Fig. 1C). We additionally recorded the TUS transducer location using neuronavigation for post hoc acoustic simulations. We also added a double-blind procedure, where both experimenter and participant were blind to which TUS condition (5Hz-rTUS vs sham) was administered, to minimise potential experimenter bias. To enable effective double-blinding, an independent researcher randomly assigned the TUS condition order across participants, using a MATLAB script. TUS parameters were automatically input into the TPO based on the defined participant and session number. Hair preparation with centrifuged ultrasound gel was initially performed after threshold and hotspot estimation, and finalised after the baseline TMS block to minimise preparation time between baseline TMS and TUS application. During both 5Hz-rTUS and sham, Gaussian white noise was played through bone-conductive headphones (Sportz3, AfterShokz, New York, USA) to maximise blinding of each condition [4]. Participants selected the maximum volume for the white noise that they found acceptable.

### Data preprocessing and analysis

Raw EMG data were exported from Signal (version 7.01; Cambridge Electronic Design, UK) to MATLAB (version 9.7.0; R2019b). Peak-to-peak MEP amplitudes for each muscle (FDI, APB, ADM) were calculated using a custom script. Data and code to reproduce the results are provided here: https://doi.org/10.17605/OSF.IO/S5AG6. Within each block, trials identified as significant outliers using Grubbs’ test (1.61%) and trials with pre-contraction (1.78%) were excluded. Precontraction was defined as the root mean square (RMS) of EMG activity in the 100ms prior to TMS exceeding the block’s average by two standard deviations. All trials with an RMS exceeding a liberal threshold of 0.045 were manually inspected (n = 10427/36000; 29%). Trials where noise in the EMG signal prevented sufficient quantification of MEP amplitude were excluded (0.55% across all muscles, 0% for FDI specifically).

Paired t-tests were conducted to assess baseline differences between 5Hz-rTUS and sham for rMT, SI_1mV,_ MEP amplitude, SICI, and ICF. The MEP amplitudes for paired-pulse measures were expressed as a ratio to the mean TS.

To assess main effects and interactions with maximal statistical power, linear mixed models (LMMs) with a maximal random effects structure were fitted using the lme4 package in R [20, 21]. Statistical significance was set at a two-tailed α=0.05 and computed with t-tests using the Satterthwaite approximation of degrees of freedom. Given the right-skewed nature of MEP amplitudes, trial-level square root corrected MEP amplitudes were used for LMMs. The time course of MEP amplitudes, SICI, and ICF was tested separately with models including TUS Condition (5Hz-rTUS/sham) and Timepoint (Baseline/T5/T30/T60) as factors. TUS-induced changes in corticospinal excitability were further tested for MEP amplitudes expressed as a ratio to the baseline mean (i.e. baseline corrected), with TUS Condition (5Hz-rTUS/sham) and Timepoint (T5/T30/T60) as factors.

In addition to LMMs, the previously implemented statistical procedures of the original study [1] were replicated using two-way rm-ANOVA on raw data without square root correction, with factors TUS Condition (5Hz-rTUS/sham) and Timepoint (Baseline/T5/T30/T60; see Supplementary Results).

### Acoustic Simulations

To determine the actual anatomical location targeted by TUS, we conducted simulations of acoustic wave propagation for each individual, using k-Plan software, a user interface for the pseudospectral time-domain solver k-Wave [22]. To generate compatible skull images, a toolbox with pre-trained deep learning convolutional neural networks was used to convert T1-weighted MRI scans to pseudo-CT images [23]. Next, the four-element CTX500 transducer location was imported from the neuronavigation TUS trajectory captured during the real 5Hz-rTUS session (exported as 3D coordinates in Brainsight coordinate space). The simulation was run using six grid points per wavelength for a single pulse duration to obtain the steady-state pressure field.

Using a custom MATLAB script, the k-Plan pressure field and grid settings were extracted using k-plan-matlab-tools (https://github.com/ucl-bug/k-plan-matlab-tools). We resampled the simulation grid to have the same spacing as the anatomical scans (i.e., T1w and pseudo-CT). Pulse average intensity was calculated from the simulated acoustic pressure amplitude p, using the plane wave approximation I= p^2/2ρc, were the density ρ was 1000 kg/m^3^ and the sound speed c was 1500 m/s.

We were primarily interested in determining the intersection between the acoustic focus and the anatomical location of the M1 hand area within the vicinity of the precentral gyrus. For this, we first segmented structural MRI scans into different tissue types, using SPM12 (https://www.fil.ion.ucl.ac.uk/spm/software/spm12/). Grey matter, white matter, and cerebrospinal fluid masks were merged to generate a binary brain mask per subject. In native space, we identified the omega formation in the pre-central gyrus at a depth of ∼30mm, and the lip of the precentral gyrus at a depth of ∼18mm [15, 24]. We extracted the location and value of the I_SPPA_ for each individual and calculated the Euclidean distance between the I_SPPA_ location and these two anatomical landmarks (Fig. 4E). We defined the acoustic focus as the volume where the intensity values are equal to or higher than half of the intensity maximum in the brain, equivalent to the full-width half-maximum intensity. Table S2 reports the simulation of free water sonication and thermal simulation results, in line with recent reporting guidelines [25]. K-Plan was also used to run a thermal simulation for one representative participant for a full 80 seconds PTD with no cooling time.

For group-level plots, the T1-weighted MRI scans and intensity maps were resampled in MNI standard space at a 1 mm isometric resolution. The binary acoustic focus maps in standard space were summed and overlaid on a standard brain with MRIcroGL. To quantify targeting accuracy through spatial overlap between the acoustic focus and the target, we required a volumetric definition of the M1 target. Therefore, we first delineated M1 with a 15 mm radius spherical region of interest (ROI) that covered both the lip of the precentral gyrus and the omega formation (Fig. S2). This ROI was chosen because stimulation of both these areas, and likely intermediate regions as well, can elicit effects of TUS on CSE [15]. We then calculated the percentage of the acoustic focus that fell within the ROI, as well as the peak ultrasound intensity within the ROI.

## Results

None of the participants reported discomfort or adverse effects following participation. Participants could not hear 5Hz-rTUS over the auditory mask, nor could they distinguish between the 5Hz-rTUS and sham sessions.

### No significant difference in baseline physiological measures

At baseline, no differences were observed for MEP amplitude (*t*(14)=0.837, *p*=.417, *d*=0.216), SICI (*t*(14)=-0.619, *p*=0.546, *d*=-0.160), or ICF (*t*(14)=-0.724, *p*=0.481, *d*=-0.187; Fig. S3). Moreover, there were no differences between baseline 5Hz-rTUS and sham conditions for rMT (t-tests; *t*(14)=0.811, *p*=0.431, *d*=0.209) or SI_1mV_ (*t*(14)=1.353, *p*=0.198, *d*=0.349; Fig. S3).

### No significant effect of 5Hz-rTUS on corticospinal or intracortical excitability

To examine the time course of corticospinal excitability changes in the primary muscle of interest (FDI), a linear mixed model was fitted predicting square root corrected MEP amplitude by Condition (5Hz-rTUS/sham), Timepoint (Baseline/T5/T30/T60) and their interaction. No significant effects were observed (Timepoint: *F*(3,14)=1.757, *p*=0.201, η□²=0.273; Condition: *F*(1,14)=1.855, *p*=0.195, η□²=0.117 Timepoint*Condition: *F*(3,14)=0.236, *p*=0.87, η□²=0.048). When expressing MEP amplitude as a ratio to baseline, we similarly found no evidence of neuromodulation (Timepoint: *F*(2,14)=1.712, *p*=0.216, η□²=0.195; Condition: *F*(1,14)=0.687, *p*=0.421, η□²=0.047; Timepoint*Condition: *F*(2,14)=0.013, *p*=0.988, η□²=0.002; Fig. 2).

**Fig. 2.**
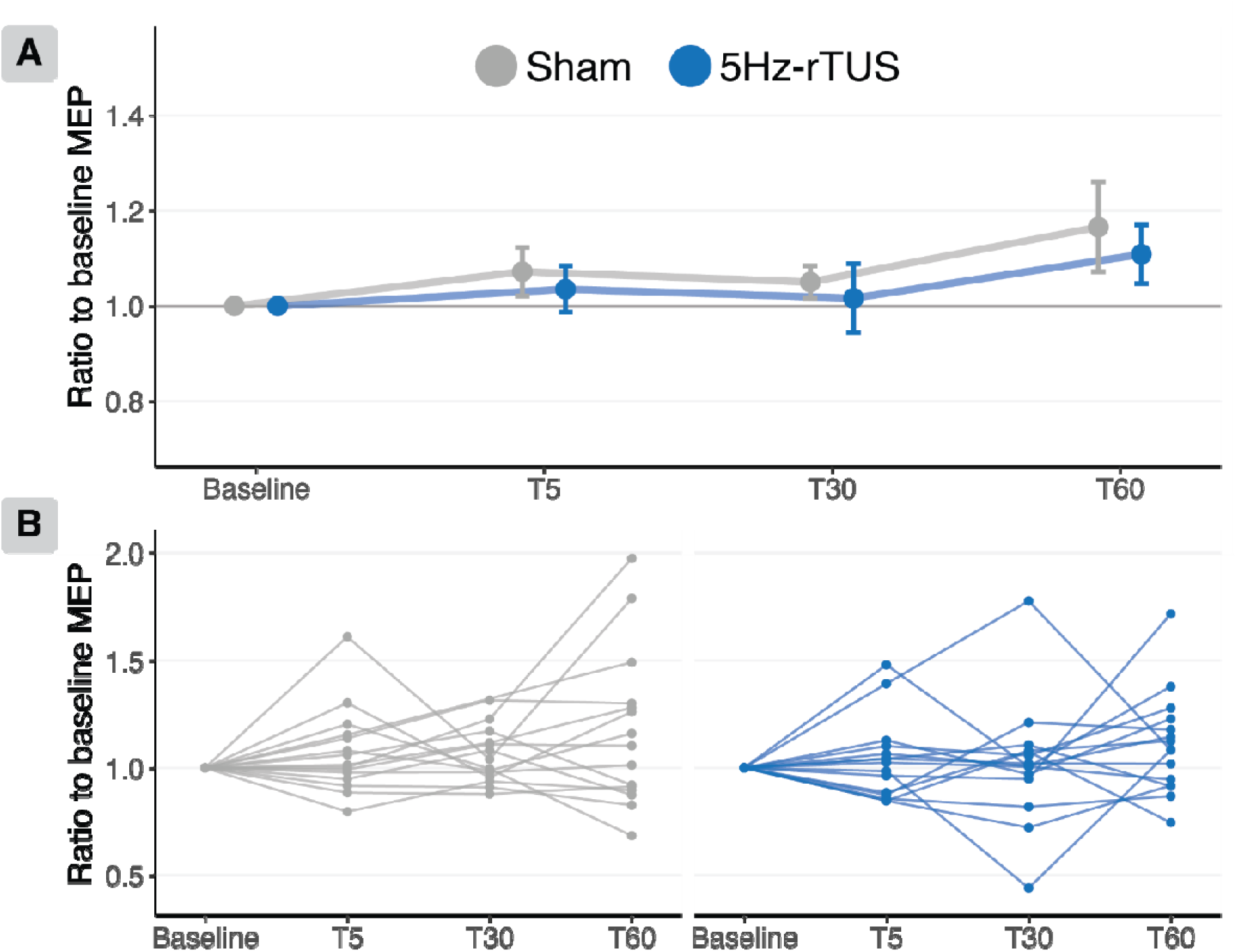
No significant effect of 5Hz-rTUS on MEP amplitude. **A.** There was no significant effect of 5Hz-rTUS on corticospinal excitability. MEP amplitudes are expressed as a ratio to baseline for each timepoint. Points and error bars represent group mean ± standard error. **B.** Participant-level data.

As shown in Fig. 3, there were also no significant effects for either SICI (Timepoint: *F*(3,14)=0.648, *p*=0.597, η□²=0.12; Condition: *F*(1, 14)=0.187, *p*=0.672, η□²=0.013; Timepoint*Condition: *F*(3,18)=0.479, *p*=0.701, η□²=0.075) or ICF (Timepoint: *F*(3,15)=0.8, *p*=0.513, η□²=0.141; Condition: *F*(1,14)=2.773, *p*=0.118, η□²=0.165 Timepoint*Condition: *F*(3,21)=0.099, *p*=0.96, η□²=0.014). RM-ANOVAs conducted on raw MEP amplitudes, following the same analyses as Zeng and co-workers, also did not reveal any significant effects [1] (see Supplementary Information 1). For the adjacent APB and ADM muscles, there was also no evidence for offline excitatory effects of sonication (Supplementary Information 2-3, Fig. S4-7). Collectively, we did not observe any reliable effect of 5Hz-rTUS on CSE or intracortical excitability.

**Fig. 3.**
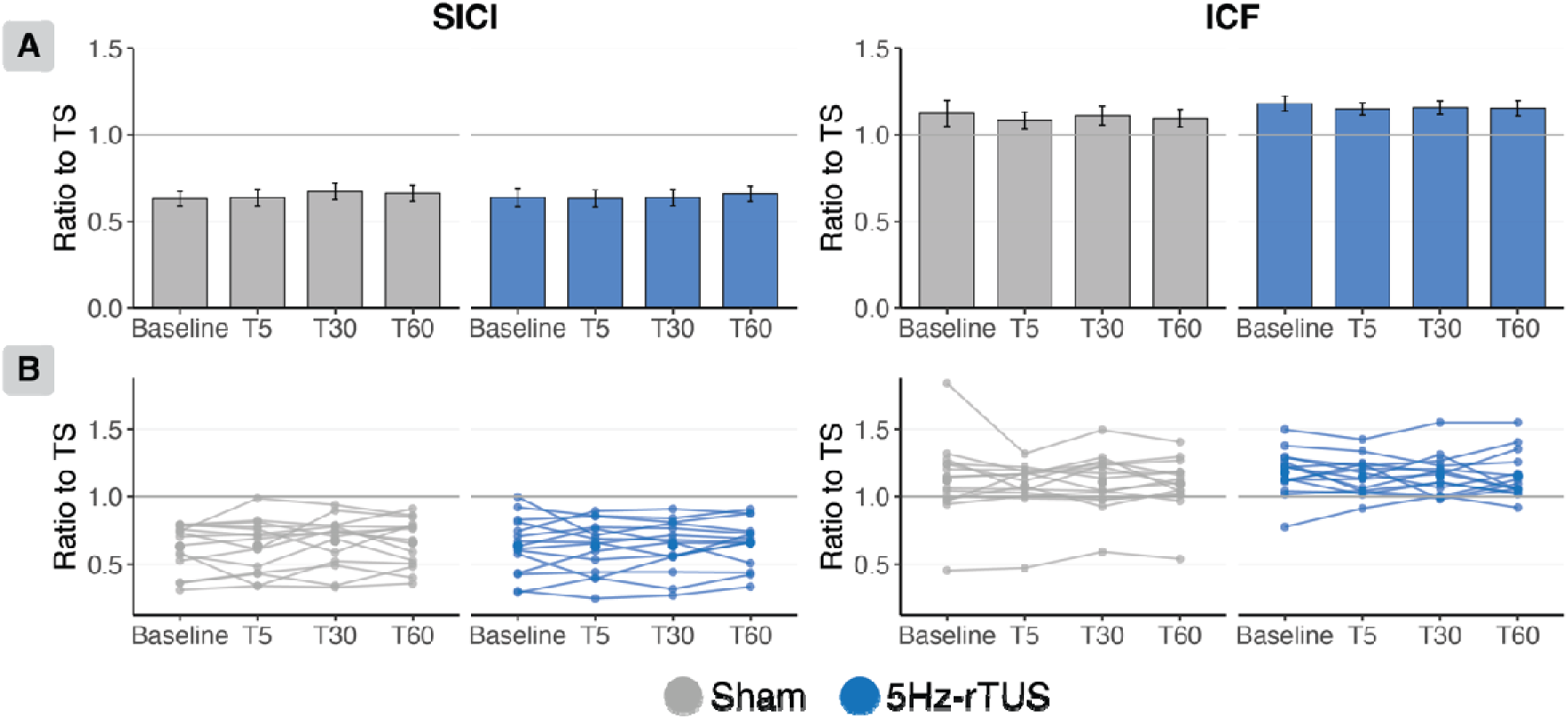
No significant effect of 5Hz-rTUS on SICI or ICF. **A.** For both SICI and ICF, there are no significant differences between sham and TUS at any time point. Data: Mean ± standard error. **B.** Participant-level data.

### No significant effect of 5Hz-rTUS on rMT or the stimulator intensity required to elicit 1mV amplitude MEPs

There was no significant effect of sonication on resting motor thresholds (Timepoint: F(3,42)=2.711, p=0.057, np2=0.162; Condition: F(1,14)=0.546, p=0.472, np2=0.038; Timepoint*Condition: F(3,42)=0.319, p=0.812, np2=0.022) nor SI1mV (Timepoint: F(1.6,22)=0.68, p=0.485, np2=0.046; Condition: F(1,14)=0.303, p=0.591, np2=0.021; Timepoint*Condition: F(2.1,29)=1.737, p=0.192, np2=0.11; Fig. S8).

### Variable ultrasound targeting of M1 based on TMS hotspot location

It is possible that ultrasound targeting of M1 based on the scalp location of the TMS hotspot is variable, and thereby may reduce the consistency and overall efficacy of 5Hz-rTUS. To address this, we conducted post-hoc simulations of the sonication target based on individual head models and assessed the degree of targeting variance in our population.

First, we observed that previous reports may have overestimated acoustic transmission, where the reported transcranial I_SPPA_ (2.26-2.93 W/cm^2^) was estimated by uniformly applying 75% attenuation from free-water I_SPPA_ (9.04-11.72 W/cm^2^; Table S1) [1, 9, 11, 13]. Here, we find a mean±sd transcranial I_SPPA_ of 1.2±0.4 W/cm^2^, corresponding to a ∼12% transmission rate, in line with empirically observed and theoretical estimations of percentage intensity transmission at *f*=500 kHz [15, 16].

Critically, our simulations reveal substantial variability in the location of the acoustic focus across participants (Fig. 4A,B). The acoustic focus overlapped with the M1 ROI in only 7 out of 15 participants. Only 33% of participants had more than 20% of the acoustic focus volume in the M1 ROI. In those 33% of subjects, the maximum intensity within the ROI was 1.0 ± 0.3 W/cm^2^ (Fig. 4B).

**Fig. 4.**
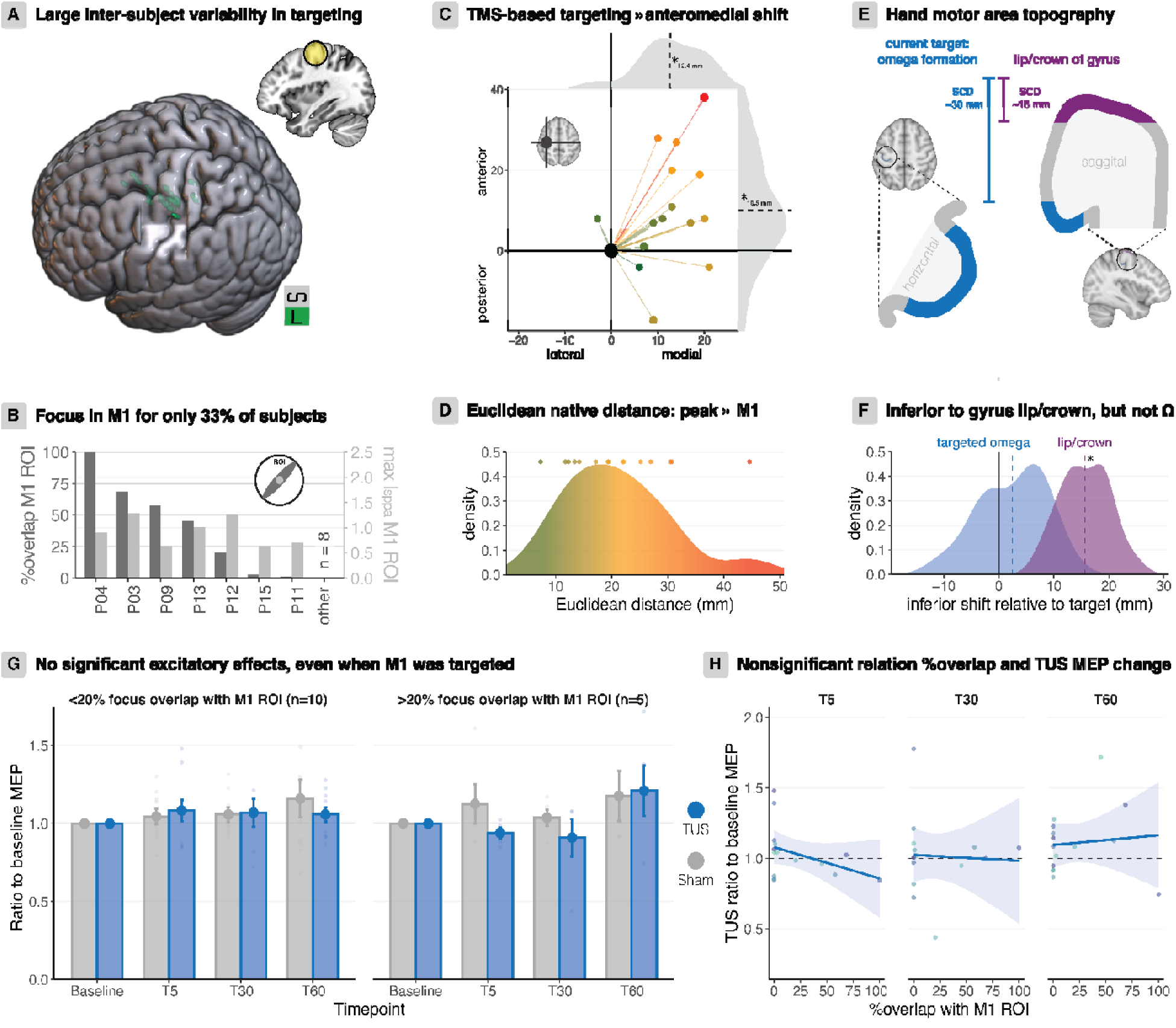
Targeting precision and offline excitatory effects of 5Hz-rTUS. **A.** Anatomical region-of-interest (15 mm M1 sphere) and variability in focus location across subjects **B.** The percentage of the acoustic focus in the M1 ROI exceeded 20% in only 5/15 participants (dark grey). Maximum ultrasound intensities within the ROI are depicted in light grey. **C.** TMS motor hotspot-based targeting leads to an anteromedial shift of the I_SPPA_. **D.** Distribution of Euclidean distances between the omega formation in the precentral gyrus and the I_SPPA_ in native space. **E.** The lip/crown of the pre-central gyrus is at a scalp-to-cortex distance of ∼18 mm, while the omega formation we targeted in this study is at a depth of ∼30 mm. **F.** There is a significant inferior shift relative to the lip/crown of the gyrus. **G.** Even when M1 was precisely targeted, there were no significant effects of sonication on corticospinal excitability. **H.** No significant association between targeting accuracy and MEP amplitude changes.

In native space, the Euclidean distance from the target omega formation seed voxel to the peak intensity voxel (I_SPPA_) was 21.1±9.5 mm (Fig. 4D). Along each 3D axis, we performed one-sample t-tests and found a significant anteromedial shift of the foci (Fig. 4C; anterior: t(14)=2.86, p=0.013; medial: t(14)=7.36, p=3.548). The degree of anterior shift (10.5±14.2 mm) and the distribution of Euclidean distances correspond with the well-known anterior shift of the TMS motor hotspot relative to the anatomical hand motor area [26–30]. We furthermore observed a significant inferior shift relative to the lip of the gyrus, which has a scalp-to-cortex distance of ∼15 mm (t(14)=14.17, p=1.079). However, there was no inferior shift relative to the targeted omega formation at a depth of ∼30 mm (t(14)=1.63, p=0.126; Fig. 4E,F). Taken together, our results demonstrate that TMS motor hotspot-based targeting method for rTUS introduces considerable dispersion around the intended target M1 location. While such misalignment between the hotspot scalp location and underlying anatomy is known, it quantifies the degree of bias and variability in directing sonication reliably to the same brain target with this procedure.

### Lack of excitatory offline 5Hz-rTUS effects when anatomical targeting is accurate

5Hz-rTUS might have been effective in participants in whom sonication was reliably directed to the presumed anatomical target region. We therefore identified those participants where the acoustic focus overlapped with the M1 ROI by >20%. However, in none of the 5 participants meeting this criterion did we observe an excitatory effect of 5Hz-rTUS on single-pulse MEP amplitude (Fig. 4G). Given the previously reported effect sizes [1], one may expect these participants to show an excitatory effect; this was not observed here. Finally, it is conceivable that there is a systematic relationship between the targeting accuracy and the sonication effect. Using linear models, we found no significant relationship between the percentage acoustic focus overlap and the ratio of post-TUS MEPs to baseline MEPs for any timepoint (Fig. 4H; T5: *b*=-0.002, *t*(14)=-1.483, *p*=0.162; T30: *b*=-4.194, *t*(14)=-0.175, *p*=0.864; T60: *b*=6.948, *t*(14)=0.342, *p*=0.738). Taken together, even when sonication is directed to M1 as intended, the excitatory effects of 5HzrTUS did not replicate.

## Discussion

Neuromodulation with TUS holds promise for clinical interventions, due to its spatial precision and potential for targeting deeper brain structures. Replication is an essential process for scientific rigour and has been earmarked by the International Transcranial Ultrasonic Stimulation Safety and Standards (ITRUSST) as crucial for accelerating TUS toward an effective neuromodulation approach. There is indeed growing attention to the rigorous experimental control required to demonstrate replicability in the field of focused ultrasound [25, 31, 32]. This need is further highlighted by examples from the fields of electrical and magnetic stimulation, where initial reports of novel stimulation protocols have often been followed by more nuanced appraisals of their efficacy [33, 34].

In the present study, we sought to replicate recently published neuromodulatory effects of offline 5Hz-rTUS directed to M1[1]. Our results did not reveal a significant effect of 5Hz-rTUS on corticospinal or intracortical excitability, contrasting with the 14 out of 15 participants showing enhanced corticospinal excitability in response to 5Hz rTUS in the original study[1], and the similarly large effect sizes of the same protocol in subsequent studies by the same group [1, 8–14].

### Variable effects of 5Hz-rTUS

The absence of significant effects held for the targeted FDI and the adjacent APB and ADM muscles. Here we initially assessed our results using linear mixed models, which have greater statistical power than rm-ANOVAs. However, our null results were the same when employing the statistical procedures as in the original study [1]. Additional qualitative assessment of individual data indicated that in only 2 out of 15 participants of the current study were the observed MEP changes consistent with the originally reported effects at T5 and T30. These results suggest the effects of 5Hz-rTUS protocol across different cohorts are likely more nuanced and less robust than originally appreciated.

This conclusion aligns with a recent replication from another group showing the opposite, i.e., inhibitory rather than excitatory effects lasting approximately 30 minutes after offline 5Hz-rTUS [15]. How the same TUS protocol directed to the same neural structure with the same overall approach can lead to completely opposite neuromodulatory effects is currently unclear. One major difference was that Bao and colleagues (2024) targeted TUS based on structural landmarks with *a priori* simulations at two focal depths - the lip/crown of the pre-central gyrus and one targeting the deeper, omega-shaped formation of the pre-central gyrus. Inhibitory effects of 5Hz-rTUS were observed at both stimulation depths.

While one cannot rule out that relatively subtle differences in applied sonication intensity between the two studies explain the opposite effects on excitability, a complete reversal of the neuromodulatory effects with such subtle intensity variation raises concerns about the clinical utility of such protocols, and certainly provides a mandate for further independent replication. Moreover, other neuromodulatory protocols used in both animal work and human research, including the 5Hz-rTUS protocol, have typically required intensities that were at least ∼300% higher (>30 W/cm^2^; 5Hz-rTUS: [35–40]. Regardless of the mechanistic explanation for how 5Hz-rTUS at very low intensities might elicit such strong neuromodulatory effects, our findings here and those by Bao and colleagues (20024) suggest that across different cohorts, the effects of low-intensity 5Hz-rTUS are more variable and potentially even orthogonal to the ones initially reported.

### Variability in targeting with TMS motor hotspot-based transducer placement

One factor that will determine the effects of TUS is the precision and consistency of its targeting. To target the primary motor cortex in combined TUS-TMS experiments, the TMS motor hotspot is commonly used as a heuristic to determine the TUS transducer location on the scalp that sonicates M1 [1, 3, 41–43]. However, this method poses obvious risks of poor target exposure to TUS.

First, when using TMS over M1, the location of the largest electrical fields and neural activation thresholds depend on several factors, including cortical folding, tissue type, and induced current direction [44, 45]. Furthermore, cortical areas closer to the lateral surface will always experience stronger electrical fields than deeper parts, due to the rapid decay of the induced electric field with increasing distance from the TMS coil [44, 45]. Therefore, the direct axial projection from the geometrical centre of the TMS coil (marked on the scalp), which is often used to determine the scalp position of the TUS transducer, is unlikely to be a reliable marker of either a specific neural structure, or the anatomical site of effective stimulation with TMS. Considering that the width of the sonication beam is considerably smaller than the spatial specificity of TMS [41, 42], even small deviations of the TUS transducer in the axial plane will lead to mis-targeting. Compared to TMS, where an altogether much larger region of the brain is targeted, this will increase the probability of directing sonication to different cortical targets across individuals.

Indeed, our post-hoc acoustic simulations demonstrate such inter-individual variability in the location of the acoustic focus relative to M1, where only 5/15 of participants showed a substantial degree of overlap between sonication and the M1 target. Scalp-based transducer placement can thus introduce inter-individual variation in the specific targeted pre- or postcentral elements. Such variation aligns with the known complexity of mapping the TMS-hotspot on the scalp to a specific anatomical target, e.g., a specific section of the omega-shaped hand knob [26–30]. Interestingly, the only study that used personalised targeting to eliminate this variability reported inhibitory effects of 5Hz-rTUS [15].

### Neuronavigation and double-blinding may explain failure to replicate

While differences in transducer model may come to mind as a possible explanation for the different findings, we would argue that the collapse from neuromodulatory effects in 14/15 participants [1] to non-significance cannot be explained solely by a difference in transducers. The axial intensity profiles between our four-element transducer and the two-element transducer used previously [1] are similar, apart from a near-field peak at 12 mm for the original two-element transducer that is too superficial to reach the brain (Fig. S1). If the neuromodulatory effects of 5Hz-rTUS were indeed entirely dependent on these very small variations in the pressure fields, it would require extraordinary precision in targeting the same neural structures consistently in every subject. This seems unlikely.

The key differences between the original work [1] and the present study were the inclusion of TMS neuronavigation and double-blinding. Using only markings on the scalp, precise TMS repositioning is challenging, particularly in terms of orientation. Without neuronavigation, slight trial-by-trial shifts in TMS position can bias MEP amplitudes [46, 47]. Further, the approach is prone to circular reasoning if the MEPs, the dependent variable, are themselves used to determine repositioning of the TMS coil. In combination with unblinded researchers, the lack of neuronavigation and double-blinding risks introduction of unconscious confirmation bias. The changes to the original work we introduced minimise such bias. It would seem prudent to suggest that inclusion of neuronavigation for TMS positioning and TUS targeting, and double-blinding should become standard ingredients of future TUS-TMS investigations.

Novel approaches to neuromodulation require realistic assessment of their efficacy, to develop methodological standards, direct further development, and ultimately help accelerate clinical use. A key component for this is independent replication. Our replication results here reappraise the effect sizes of excitatory neuromodulation of 5Hz-rTMS to M1 previously reported, but also suggest avenues toward consistent and reproducible evaluation of novel TUS protocols.

## Data Statement

Data and code to reproduce the results reported in this study are available at: https://doi.org/10.17605/OSF.IO/S5AG6

## Acknowledgements

We thank Paul Hammond for their help and assistance.

## Funding

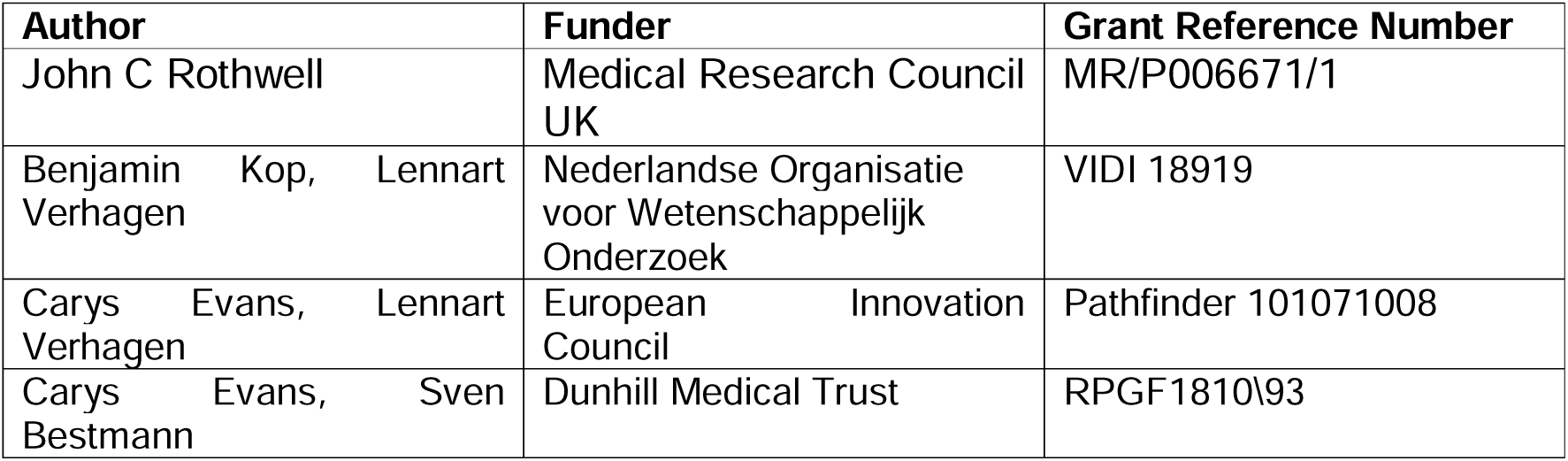

## Contributions

Po-Yu Fong: investigation, conceptualisation, methodology, data curation, writing – original draft, writing – review & editing, Resources, Supervision (the investigation team).

Benjamin Kop: conceptualisation, methodology, software, formal analysis, data curation, writing – original draft, writing – review & editing, visualisation.

Carys Evans: conceptualisation, methodology, writing – original draft, writing – review & editing, visualisation.

Vidya Gani Wijaya: investigation

Yongling Lin: investigation

Drew Cappotto: investigation

Jenny S. A. Lee: investigation

Anna Latorre: investigation

Joy Song: investigation

Bradley Treeby: writing – Resources, review & editing

Eleanor Martin: writing – Resources, review & editing

John Rothwell: Funding acquisition, Resources, Supervision

Lennart Verhagen: Methodology, Writing - review & editing, Funding acquisition, Conceptualisation, Supervision

Sven Bestmann: Supervision, conceptualisation, methodology, writing – original draft, writing – review & editing

