## Supplementary Materials for "A Double-Blind Replication Attempt of Offline 5Hz-rTUS-Induced Corticospinal Excitability"

### Supplementary Material

#### Comparison of axial intensity profile between present and prior work

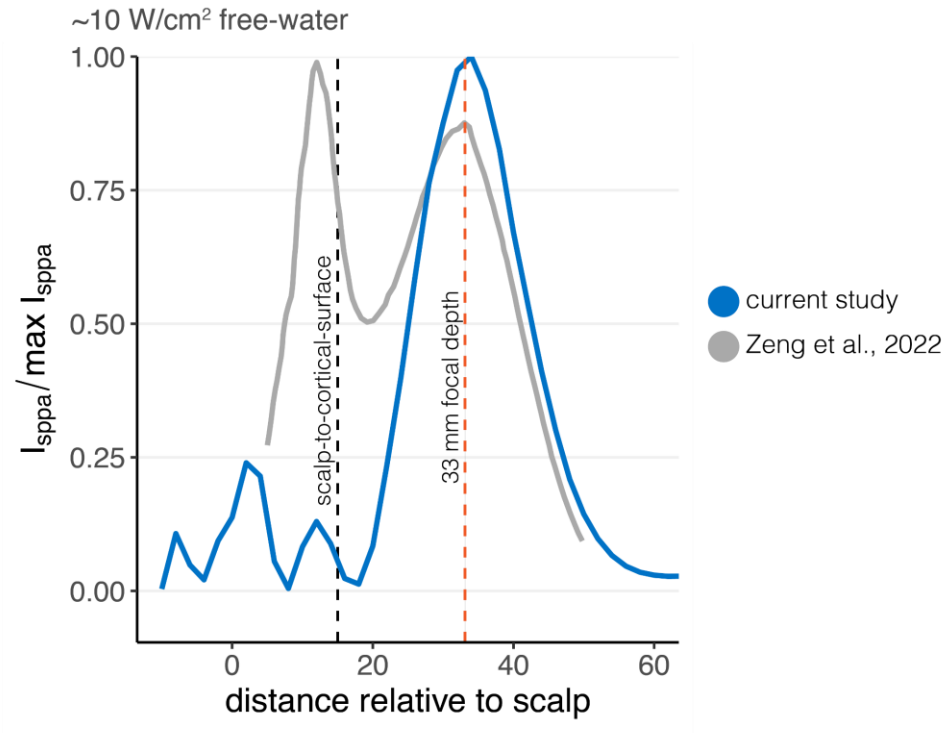


**Supplementary Fig. 1. Comparison of normalised axial intensity profiles**

The peak intensity of the intended focus using transducer H246 was 33 mm for Zeng and colleagues (2022; dashed orange line). We replicate the intended focal depth of Zeng et al (2022) by using a TPO focal depth setting of 43.5 mm for CTX500-025, and accounting for the use of a 10 mm gel pad. We show a similar axial profile. An average scalp-to-cortical-surface distance of 15 mm is depicted with a dashed black line[1]. Plots adapted from the original report[2].

For the present study, the axial profiles for CTX500-025 measured by the manufacturer (Sonic Concepts, WA, USA) at similar focal depths (grey) were interpolated to estimate the pressure distribution for the currently applied focal depth of 43.5 mm (blue). The normalised axial profile data displayed in the original study [2] were extracted using WebPlotDigitizer (version 5; CA, USA) by digitising the axes, performing automatic extraction using a pen tool and a colour filter, and sampling at a 5-pixel interval.

**Supplementary Table 1**

| | **Transducer** | **Power** | **$I_{sppa\_fw}$** | **$I_{sppa\_tc}$** | **Simulated $I_{sppa\_tc}$** |
|---|---|---|---|---|---|
| Zeng et al., 2022 | H246† | 20W* | 9.04 W/cm$^2$ | 2.26** W/cm$^2$ | - |
| Present study | CTX500-025† | 3.71W* | 10.0 W/cm$^2$ | 2.50** W/cm$^2$ | 1.20 ± 0.43 W/cm$^2$ |

†Transducers from Sonic Concepts (Bothell, WA, USA).
*****Power levels required to reach an equivalent $I_{sppa}$ differ for different transducers. Higher power does not suggest that 'more stimulation' was applied.
**Transcranial $I_{sppa}$ ($I_{sppa\_tc}$) is estimated by Zeng and colleagues [2] by applying 75% attenuation from free-water $I_{sppa}$ ($I_{sppa\_fw}$).

**Supplementary Table 2**

**Transducer and Drive System Parameters**

| | Manufacturer, Model Number | Centre Frequency | Radius of curvature | Aperture Diameter | Number of Elements | Element Distribution |
|---|---|---|---|---|---|---|
| Transducer | Sonic Concepts, CTX-500-025 | 500 kHz | 64 mm | 64 mm | 4 | Spherical cap, annular array, equal area, |
| Matching | 4 channel electrical impedance matching network, Sonic Concepts | | | | | |
| Drive system | NeuroFUS Pro 4 channel TPO | | | | | |

**Driving System Settings**

| | Operating Frequency | Output level Setting | Focal Position Setting |
|---|---|---|---|
| Motor .. | 500 kHz | 10 W/cm$^2$ | 43.5 mm |

**Free Field Pressure Parameters**

| | Spatial Peak Pressure Amplitude | Position of Spatial Peak Pressure | Axial Focal Size (-3dB) | Lateral Focal Size (-3dB) | Axial Focal Size (-6dB) | Lateral Focal Size (-6dB) |
|---|---|---|---|---|---|---|
| 43.5 mm | 548 ± kPa | [0, 0, 53.5] mm | 14.2 mm | 2.6 mm | 19.7 mm | 3.5 mm |

Notes: Reference position is at centre of the spherical array surface.

28

**Pulse Timing Parameters**

| | | Duration | Ramp Duration | Ramp Shape | Repetition Interval / Frequency |
|---|---|---|---|---|---|
| Experiment 2 | Pulse | 20 ms | 0 ms | No ramp | 0.2 s / 5 Hz |
| | Pulse Train | 80 s | | | |

Notes:

**In Situ Exposure Parameters**

| | Spatial Peak Pressure Amplitude [kPa] | Spatial peak pulse average intensity [W/cm2] | Mean position of Spatial Peak Pressure [mm] | Axial Focal Size (-3dB) [mm] | Lateral Focal Size (-3dB) [mm] | Axial Focal Size (-6dB) [mm] |
|---|---|---|---|---|---|---|
| Motor | 189 ± 39 | 1.2 ± 0.43 | [0.3 ± 0.5, -0.1 ± 0.5, 51.1 ± 1.3] mm | 17.0 ± 3.1 | [3.1 ± 0.2, 2.9 ± 0.1] | 32.9 ± 14.3 |

Notes: Reference position is at centre (origin) of spherical array surface

| Mechanical Index | Maximum temperature rise |
| --- | --- |
| $0.26 \pm 0.05$ | 0.22 °C |

32

#### M1 ROI

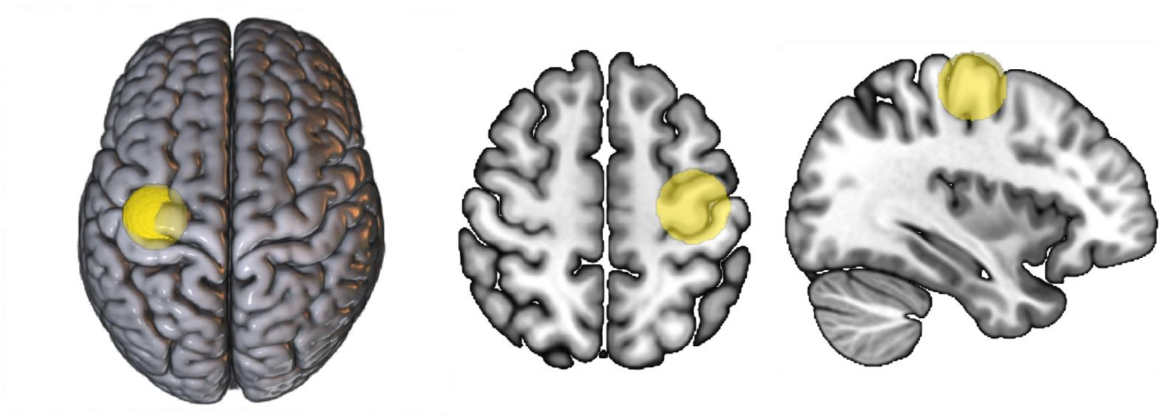

**Supplementary Fig. 2**. 15 mm M1 spherical region-of-interest (ROI). The ROI was structurally identified in MNI152 space and used to assess targeting. The ROI was placed such that the both the lip/crown of the precentral gyrus and the omega formation are included.

#### No baseline differences between sham and 5Hz-rTUS

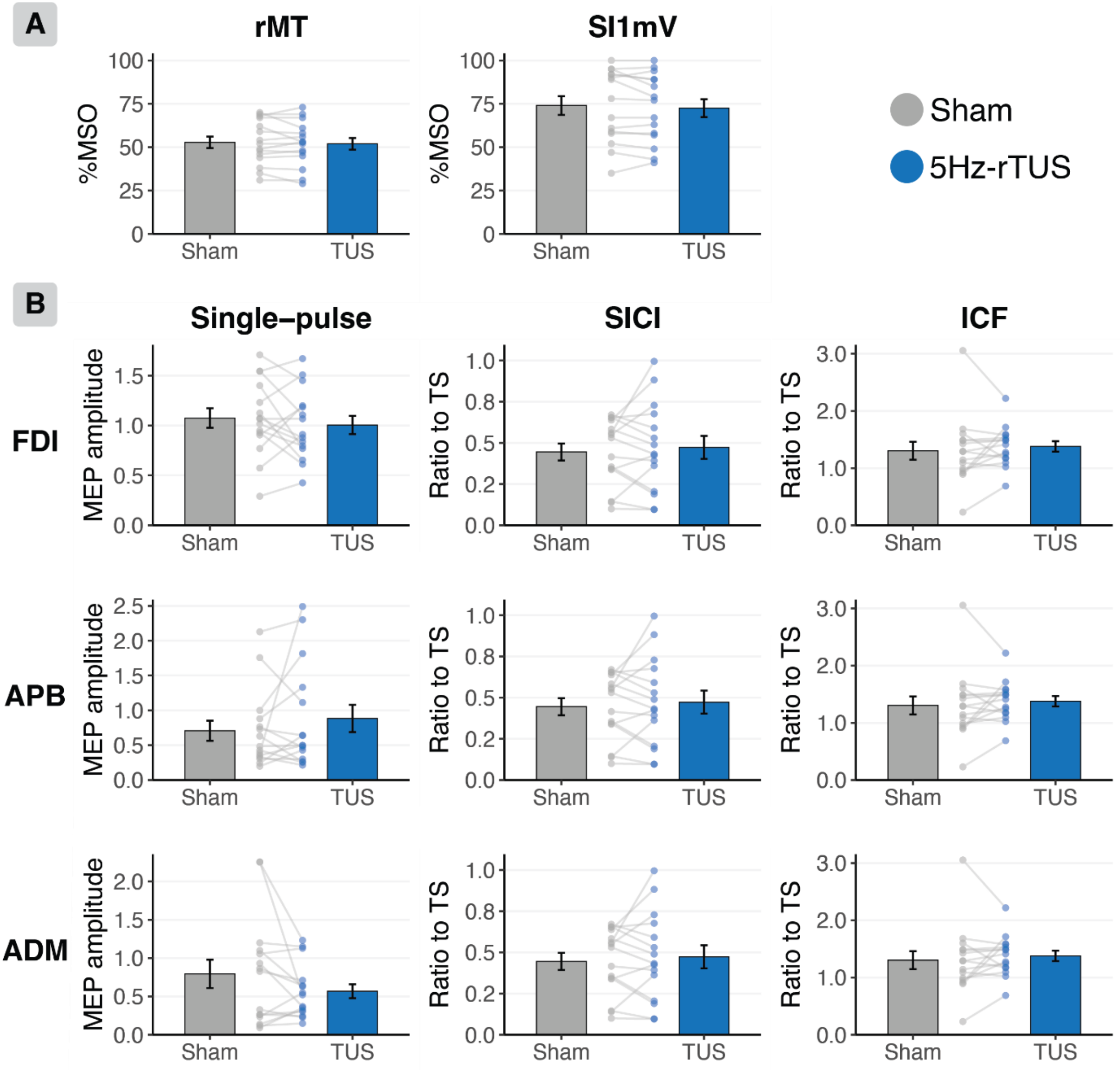


**Supplementary Fig. 3. No significant differences between sham and 5Hz-rTUS at baseline**

**A.** Sham vs 5Hz-rTUS effects on resting motor threshold (rMT) and the TMS intensity required to evoke a ~1 mV MEP ($SI_{1mV}$).

**B.** Single-pulse MEP amplitude (left), short interval cortical inhibition (SICI; middle), and intracortical facilitation (ICF; right) measured from the FDI (top), APB (middle), or ADM (bottom) muscles, respectively.

### Supplementary Information 1

#### Replicated statistical methodology (rm-ANOVAs)

Repeated measures ANOVAs (rm-ANOVA) on raw MEP amplitudes were conducted to replicate the statistical methods employed by Zeng and colleagues[2]. To examine the time course of corticospinal excitability changes in MEP amplitudes for the primary muscle of interest, the FDI, a rm-ANOVA with factors Condition (5Hz-rTUS/sham), Timepoint (Baseline/T5/T30/T60), and their interaction was conducted. This analysis revealed a trend for Timepoint ($F(1.77,23) = 2.967$, $p = 0.077$, $\eta_p^2 = 0.186$), but no significant effect of Condition ($F(1,13) = 1.903$, $p = 0.191$, $\eta_p^2 = 0.128$), nor a significant Condition*Timepoint interaction ($F(1.57,20) = 0.284$, $p = 0.703$, $\eta_p^2 = 0.021$). To further examine TUS-induced changes in corticospinal excitability, MEP amplitudes were expressed as a ratio to baseline and tested with the factors Condition (5Hz-rTUS), Timepoint (T5/T30/T60), and their interaction. No significant effects were observed (Timepoint: $F(1.33,17) = 2.465$, $p = 0.128$, $\eta_p^2 = 0.159$; Condition: $F(1,13) = 0.204$, $p = 0.659$, $\eta_p^2 = 0.015$; Timepoint*Condition: $F(2,26) = 0.045$, $p = 0.956$, $\eta_p^2 = 0.003$). In line with these findings, no significant effects were observed for paired-pulse measures (SICI: Timepoint: $F(3,42) = 0.463$, $p = 0.71$, $\eta_p^2 = 0.032$; Condition: $F(1,14) = 0.008$, $p = 0.932$, $\eta_p^2 = 0.001$; Timepoint*Condition: $F(3,42) = 0.819$, $p = 0.491$, $\eta_p^2 = 0.055$; ICF: Timepoint: $F(3,42) = 1.107$, $p = 0.357$, $\eta_p^2 = 0.073$; Condition: $F(1,14) = 2.283$, $p = 0.153$, $\eta_p^2 = 0.14$; Timepoint*Condition: $F(1.76,25) = 0.055$, $p = 0.929$, $\eta_p^2 = 0.004$). Taken together, results from closely replicated analyses do not provide evidence for effective ultrasonic neuromodulation of corticospinal excitability, in line with the linear mixed models used in the main text.

### Supplementary Information 2

#### Results for abductor pollicis brevis (APB)

A linear mixed model predicting square root corrected MEP amplitude measured over the APB by Condition (5Hz-rTUS/sham), Timepoint (Baseline/T5/T30/T60) and their interaction revealed a significant effect of Timepoint ($F(3,14) = 4.618$, $p = 0.019$, $\eta_p^2 = 0.498$), with MEP amplitudes increasing over time. However, no significant effect of Condition ($F(1,14) = 0.123$, $p = 0.731$, $\eta_p^2 = 0.009$) or Timepoint*Condition ($F(3,14) = 1.102$, $p = 0.381$, $\eta_p^2 = 0.191$) was observed. When testing MEP amplitude expressed as a ratio to baseline, no significant effects were observed (Timepoint: $F(2,14) = 1.931$, $p = 0.182$, $\eta_p^2 = 0.219$; Condition: $F(1,14) = 2.553$, $p = 0.132$, $\eta_p^2 = 0.154$; Timepoint*Condition: $F(2, 14) = 0.697$, $p = 0.515$, $\eta_p^2 = 0.094$). A significant effect of Condition on SICI was observed (Condition: $F(1,14) = 6.338$, $p = 0.024$, $\eta_p^2 = 0.309$), with SICI being more pronounced for 5Hz-rTUS (i.e., lower ratio) compared to sham. However, there was no significant interaction with Timepoint ($F(3,16) = 2.134$, $p = 0.136$, $\eta_p^2 = 0.288$), nor a main effect thereof ($F(3,17) = 0.971$, $p = 0.429$, $\eta_p^2 = 0.144$). No significant effects were observed for ICF (Timepoint: $F(3,14) = 0.916$, $p = 0.458$, $\eta_p^2 = 0.16$; Condition: $F(1,14) = 2.872$, $p = 0.112$, $\eta_p^2 = 0.17$ Timepoint*Condition: $F(3,16) = 0.856$, $p = 0.483$, $\eta_p^2 = 0.135$). Taken together, these results do not provide evidence for effective ultrasonic neuromodulation of corticospinal excitability.

RM-ANOVAs similarly did not reveal evidence for effective ultrasonic neuromodulation (*Raw MEP amplitude*: Timepoint: $F(3,39) = 3.133$, $p = 0.036$, $\eta_p^2 = 0.194$; Condition: $F(1,13) = 0.061$, $p = 0.808$, $\eta_p^2 = 0.005$; Timepoint*Condition: $F(3,39) = 1.445$, $p = 0.245$, $\eta_p^2 = 0.1$; *MEP amplitude as ratio to baseline*: Timepoint: $F(2,26) = 1.382$, $p = 0.269$, $\eta_p^2 = 0.096$; Condition: $F(1,13) = 1.489$, $p = 0.244$, $\eta_p^2 = 0.103$; Timepoint*Condition: $F(2,26) = 0.472$, $p = 0.629$, $\eta_p^2 = 0.035$; *SICI*: Timepoint: $F(3,42) = 3.185$, $p = 0.033$, $\eta_p^2 = 0.185$; Condition: $F(1,14) = 1.308$, $p = 0.272$, $\eta_p^2 = 0.085$; Timepoint*Condition: $F(1.66,23) = 0.439$, $p = 0.614$, $\eta_p^2 = 0.03$; *ICF*: Timepoint: $F(1.85,26) = 0.714$, $p = 0.489$, $\eta_p^2 = 0.049$; Condition: $F(1,14) = 0.49$, $p = 0.496$, $\eta_p^2 = 0.034$; Timepoint*Condition: $F(2.09,29) = 0.583$, $p = 0.572$, $\eta_p^2 = 0.04$).

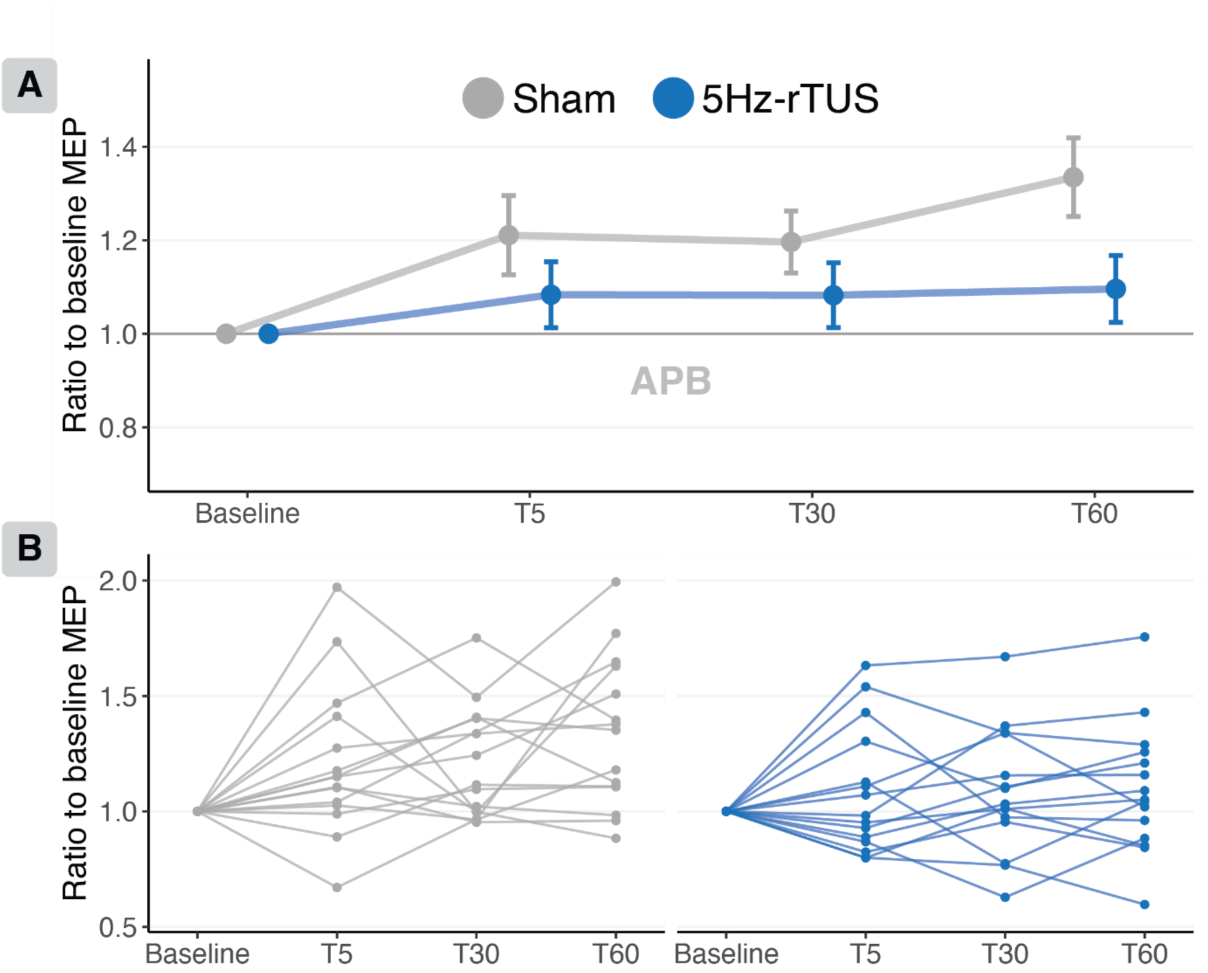


**Supplementary Fig. 4. No significant effect of TUS on MEP amplitude measured over the adjacent abductor pollicis brevis (APB).**

- **A.** There was a significant main effect of Timepoint (Baseline/T0/T5/T30/T60), but no significant main effect or interaction with Condition (5Hz-rTUS/sham). MEP amplitudes are expressed as a ratio to baseline for each timepoint. Heavyweight points: Group mean ± standard error.
- **B.** Participant-level data.

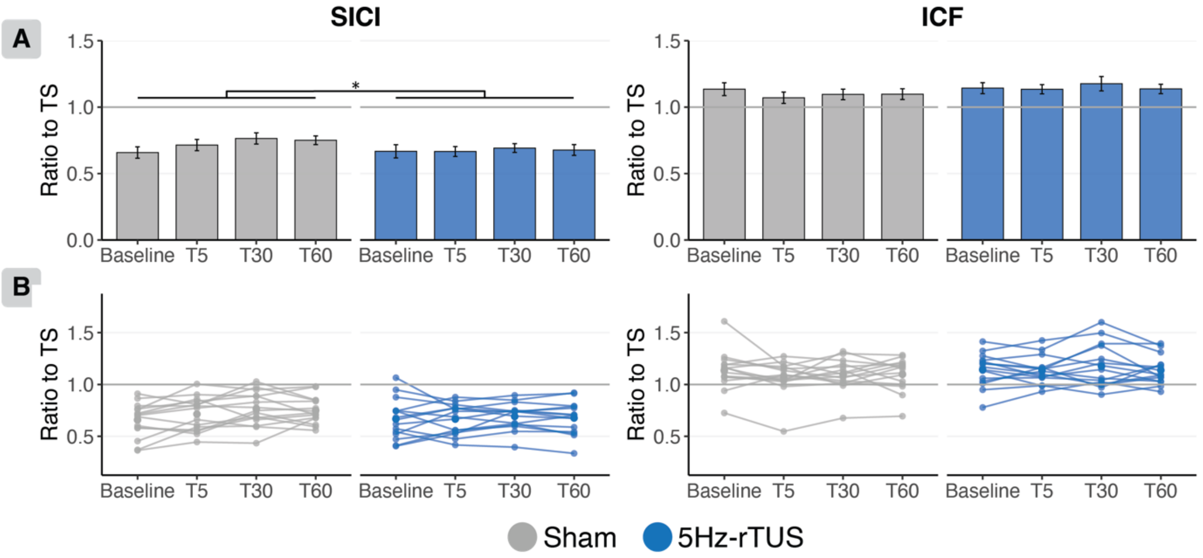


**Supplementary Fig. 5. No significant excitatory effect of TUS on SICI or ICF for APB.**

**A.** For SICI there was a significant effect of Condition (5Hz-rTUS/sham), but SICI was more pronounced for 5Hz-rTUS than sham, pointing towards inhibition rather than excitation. For ICF, there was no significant effect of TUS. MEP amplitudes are expressed as a ratio to the test-stimulus. Data and error bars represent group mean ± standard error.

**B.** Participant-level data.

#### Supplementary Information 3

#### Results for abductor digiti minimi (ADM)

A linear mixed model predicting square root corrected MEP amplitude measured over the ADM by Condition (5Hz-rTUS/sham), Timepoint (Baseline/T5/T30/T60) and their interaction revealed a significant effect of Timepoint ($F(3,14) = 4.674$, $p = 0.018$, $\eta_p^2 = 0.493$), with MEP amplitude increasing over time. No significant effect of Condition ($F(1,14) = 1.582$, $p = 0.229$, $\eta_p^2 = 0.101$) or Timepoint*Condition ($F(3, 14) = 1.396$, $p = 0.285$, $\eta_p^2 = 0.23$) was observed. When testing MEP amplitude expressed as a ratio to baseline, no significant effects were observed (Timepoint: $F(2,14) = 1.548$, $p = 0.247$, $\eta_p^2 = 0.179$; Condition: $F(1,14) = 0.874$, $p = 0.366$, $\eta_p^2 = 0.059$; Timepoint*Condition: $F(2, 14) = 2.118$, $p = 0.157$, $\eta_p^2 = 0.231$). Similarly, no significant effects were observed for either SICI or ICF (SICI: Timepoint: $F(3,16) = 2.219$, $p = 0.126$, $\eta_p^2 = 0.298$; Condition: $F(1,14) = 0.306$, $p = 0.589$, $\eta_p^2 = 0.022$; Timepoint*Condition: $F(3,16) = 0.04$, $p = 0.989$, $\eta_p^2 = 0.007$; ICF: Timepoint: $F(3,15) = 0.968$, $p = 0.434$, $\eta_p^2 = 0.165$; Condition: $F(1,14) = 1.875$, $p = 0.193$, $\eta_p^2 = 0.118$; Timepoint*Condition: $F(3,17) = 1.739$, $p = 0.196$, $\eta_p^2 = 0.231$).

RM-ANOVAs similarly did not reveal evidence for effective ultrasonic neuromodulation (*Raw MEP amplitude*: Timepoint: $F(1.45,19) = 2.948$, $p = 0.09$, $\eta_p^2 = 0.185$; Condition: $F(1,13) = 1.971$, $p = 0.184$, $\eta_p^2 = 0.132$; Timepoint*Condition: $F(1.58,21) = 1.834$, $p = 0.189$, $\eta_p^2 = 0.124$; *MEP amplitude as ratio to baseline*: Timepoint: $F(2,26) = 0.775$, $p = 0.471$, $\eta_p^2 = 0.056$; Condition: $F(1,13) = 0.352$, $p = 0.563$, $\eta_p^2 = 0.026$; Timepoint*Condition: $F(2,26) = 1.329$, $p = 0.282$, $\eta_p^2 = 0.093$; *SICI*: Timepoint: $F(1.68,23) = 0.539$, $p = 0.56$, $\eta_p^2 = 0.037$; Condition: $F(1,14) = 0.035$, $p = 0.854$, $\eta_p^2 = 0.003$; Timepoint*Condition: $F(1.84,26) = 0.657$, $p = 0.514$, $\eta_p^2 = 0.045$; *ICF*: Timepoint: $F(3,42) = 1.481$, $p = 0.234$, $\eta_p^2 = 0.096$; Condition: $F(1,14) = 0.368$, $p = 0.554$, $\eta_p^2 = 0.026$; Timepoint*Condition: $F(3,42) = 0.425$, $p = 0.736$, $\eta_p^2 = 0.029$).

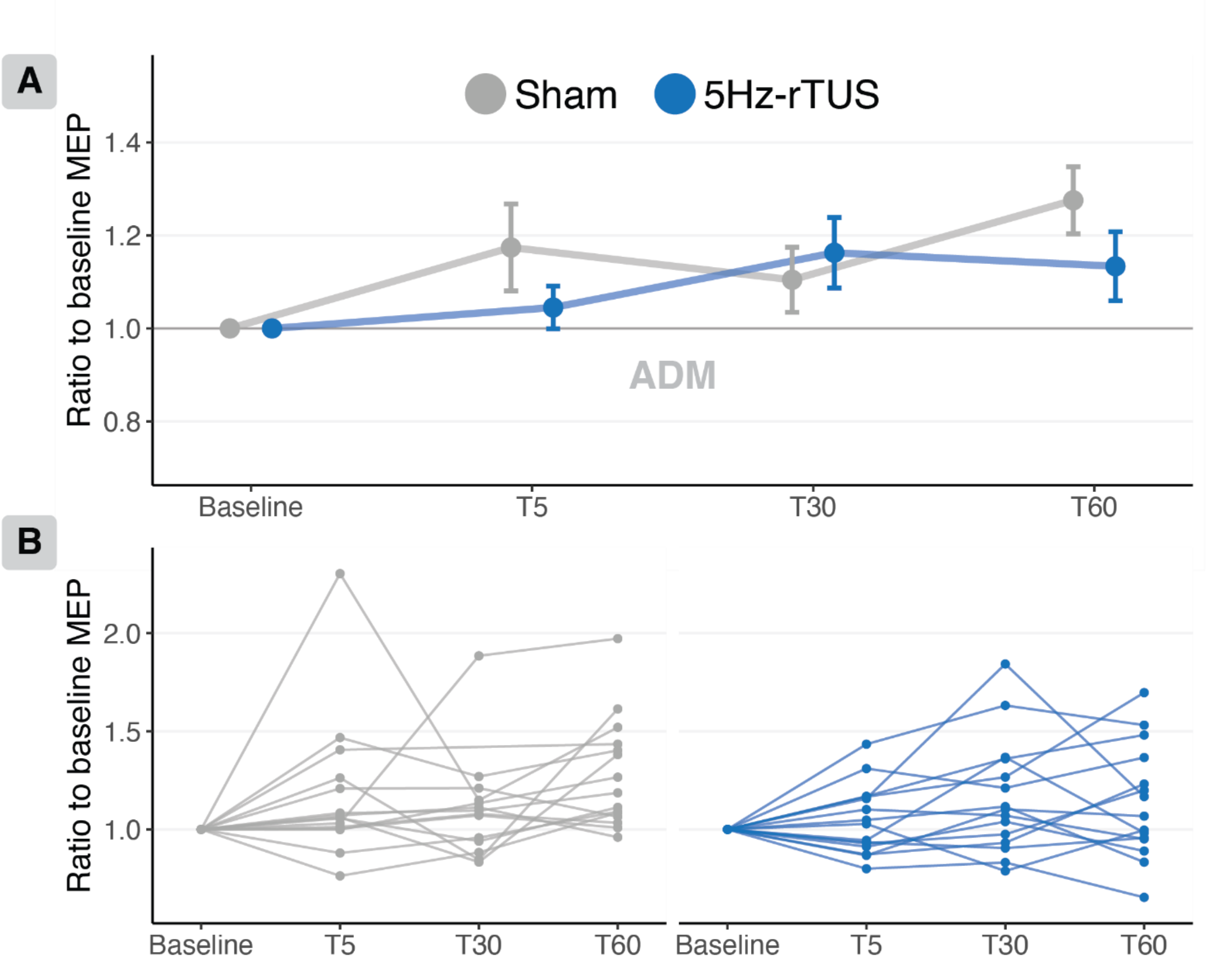


**Supplementary Fig. 6. No significant effect of 5Hz-rTUS/sham on MEP amplitude measured over the adjacent abductor digiti minimi (ADM).**

**A.** There was a significant main effect of Timepoint (Baseline/T0/T5/T30/T60), but no significant main effect or interaction with Condition (5Hz-rTUS/sham). MEP amplitudes expressed as a ratio to baseline for each timepoint. Points and error bars represent group mean ± standard error.

**B.** Participant-level data.

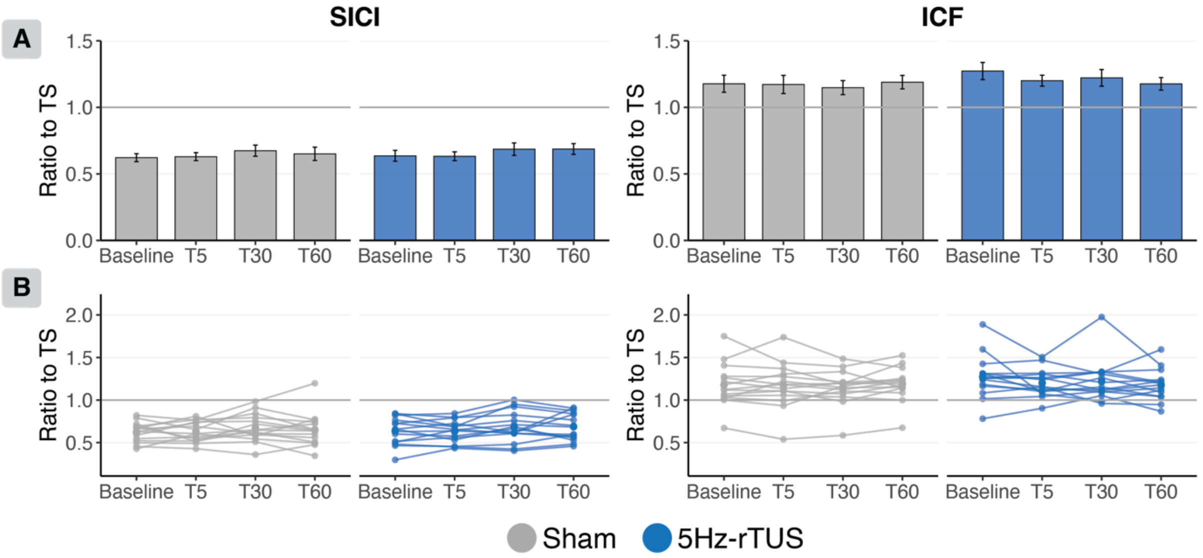


**Supplementary Fig. 7. No significant effect of 5Hz-rTUS/sham on SICI or ICF. MEP amplitudes are expressed as a ratio to the test-stimulus.**

- **A.** For both SICI and ICF, there are no significant differences between sham and 5Hz-rTUS at any time point. Data and error bars represent group mean ± standard error.
- **B.** Participant-level data.

#### No significant effect of 5Hz-rTUS on resting motor threshold or SI1mV

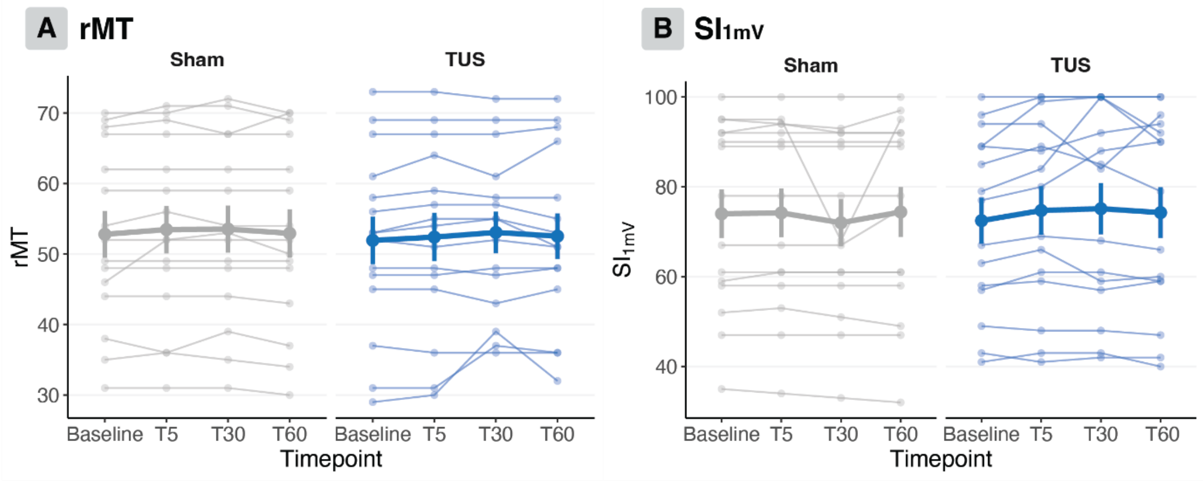


**Supplementary Fig. 8. No significant effect of 5Hz-rTUS on resting motor threshold or SI1mV**

- **A.** No significant effect of 5Hz-rTUS on resting motor threshold (rMT). Data: Mean ± standard error. Participant-level data: smaller points and lines.
- **B.** No significant effect of 5Hz-rTUS on the stimulator intensity required to evoke a ~1mV MEP ($SI_{1mV}$). Heavyweight points and error bars represent group mean ± standard error. Lightweight points represent participant-level data.
